# Homeostatic Activation of 26S Proteasomes by Protein Kinase A Protects against Cardiac and Neurobehavior Malfunction in Alzheimer’s Disease Mice

**DOI:** 10.1101/2025.03.28.645869

**Authors:** Anand Chakroborty, Saima Ejaz, Jack O. Sternburg, Yasin Asadi, Mingqi Cai, Abena Asua Dwamena, Samiksha Giri, Oluwagbemisola Adeniji, Md Salim Ahammed, Erin Abigail Gilstrap, Md Giash Uddin, Caroline McDowell, Jinbao Liu, Hongmin Wang, Xuejun Wang

**Author notes:** Adress correspondence to Xuejun Wang, MD, PhD, University of South Dakota Sanford School of Medicine, Vermillion, SD 57069, USA; or Hongmin Wang, PhD, TTUHSC School of Medicine, Lubbock, TX 79430, USA. These authors contributed equally (Anand Chakroborty, Saima Ejaz).

## Abstract

Alzheimer’s Disease (AD) patients often show brain and cardiac malfunction. AD represents a leading cause of morbidity and mortality worldwide, but the demand for effective treatment for AD is far from being met. This is primarily because AD pathogenesis, including brain-heart interaction, is poorly understood. Proteasome functional insufficiency is implicated in AD; as such, proteasome enhancement promises a potentially new strategy to treat AD. The proteasome can be activated by protein kinase A (PKA) via selectively phosphorylating Ser14-RPN6/PSMD11 (p-S14-RPN6); however, whether p-S14-RPN6 is altered and what role p-S14-RPN6 plays in AD remain unclear. Hence, this study was conducted to address these critical gaps. We found that genetic blockade of the homeostatic p-S14-Rpn6 via germline knock-in of Rpn6^S14A^ (referred to as S14A) significantly reduced proteasome activities in the cerebral cortex but did not discernibly impair learning and memory function in 4-month-old mice or cause cardiac dysfunction before 12 months of age. Increases in Ser14-phosphorylated Rpn6 in the cerebral cortex and markedly elevated Aβ proteins in the myocardium were observed in young 5XFAD mice, a commonly used AD model. When introduced into the 5XFAD mice, S14A significantly aggravated the learning and memory deficits as revealed by the radial arm water maze tests and accelerated cardiac malfunction as measured by serial echocardiography in the same cohort of 5XFAD mice. Thus, the present study establishes for the first time that homeostatic activation of 26S proteasomes by basal p-S14-RPN6 or PKA activity protects against both the brain and heart malfunction in the 5XFAD mice.

## INTRODUCTION

Characterized by a chronic dementia in the elderly, Alzheimer’s disease (AD) is an progressive neurodegenerative disorder that affects approximately 55 million people worldwide^1^. The current annual global cost of dementia is approaching US$ 1.3 trillion and is expected to rise to US$ 2.8 trillion by 2030^2^. The major segment of the population diagnosed with AD are elderlies (>65 years or above) suffering from late-onset AD (LOAD) and the remaining 5% of cases (<65 years) are surviving with distressing early-onset AD (EOAD) ^3, 4^. The estimated burden of LOAD is around 6.9 million Americans, and this number is projected to reach 13.8 million by 2060 ^5^.

AD is clinically diagnosed as the persistent decline of behavioral and cognitive functions and pathologically featured by the presence of amyloid β-protein (Aβ) plaques and the neurofibrillary tangles of either mutated or hyperphosphorylated tau protein in the brain ^6–13^. While no preventive strategy or medication exists, treatment options are limited with the recent induction of FDA-approved aducanumab, lecanemab, and donanemab to attenuate the progression during the early stages of AD ^12, 14–16^.

Nevertheless, reversal or blockage of the cognitive decline due to neurodegeneration is still far-fetched.

The ubiquitin-proteasome system (UPS) is essential in eukaryotic organisms for degradation of misfolded or abnormal proteins and for maintaining proteostasis ^17, 18^. The UPS also prevents aggregation of pathognomonic proteins within the neuron via timely removal of them and dysfunction in the pathway due to aging causes neurodegenerative diseases, including AD ^19–27^. The proteolytic machine of the UPS is the 26S proteasome complex that serves in the cytoplasm and nucleus of all eukaryotic cells to break down ubiquitinated proteins ^28–30^. The 26S holo-complex is a “dumb-shell-shaped” symmetric structure composed of barrel-shaped 20S proteasome core catalytic complex (CP) flanked at one or both ends by the 19S regulatory complex (RP) (19S-20S or 19S-20S-19S). The 19S RP consists of the proximal base and distal lid subcomplex ^18, 31^. Several studies have shown that 19S RP subunits like Rpn6, Rpn10, Rpt2, and Rpt6 are vital for proteasome activity and their diminution in the cell system triggers central aging and neurodegeneration ^24, 32–36^.

PSMD11 (proteasome subunit D11) is a mammalian orthologue of yeast Rpn6 (regulatory particle non-ATPase 6). Studies using *Caenorhabditis elegans* have revealed that RPN6/PSMD11 is a potent factor that confers resistance to proteotoxic stress and extends the lifespan of the nematode ^32, 37, 38^. The Rpn6/PSMD11 is one of the 19 lid subunits of the 19S RP and consists of an α-solenoid-like fold and a proteasome COP9/signalosome eIF3 (PCI) module in a right-handed super-helical configuration, and is stated to play a critical role in structural integrity by holding RP and CP subcomplexes together ^39, 40^. RPN6 is a critical site for the phosphorylation by the cAMP-dependent kinase (protein kinase A, PKA). PKA specifically phosphorylates RPN6 at Ser14 and thereby activates the 26S proteasome ^38, 41, 42^; both in vitro and in vivo studies have established that Ser14-phophorylated RPN6 mediates the activation of 26S proteasomes by PKA ^43^.

Although proteasome enhancement was suggested to underlie the protective effects of cAMP augmentation on AD ^43^, the role of homeostatic Ser14-RPN6 phosphorylation (p-S14-RPN6) or, by extension, the role of PKA-induced activation of 26S proteasomes in AD pathogenesis and progression has not been genetically interrogated or established. Hence, the present study was conducted to primarily address this critical gap. Taking advantage of our recently created and validated Rpn6^S14A^ knock-in mouse model (referred to as **S14A** hereafter), the present study investigated the impact of genetic blockade of p-S14-RPN6 on neurobehavior and cardiac functions in mice at baseline and, more importantly, on the progression of brain and cardiac malfunction in the 5XFAD mice, a widely used animal model of familial AD, containing 3 APP mutations and 2 PSEN1 mutations ^44^. Our results indicate that genetic blockade of p-S14-Rpn6 exerts rather modest effects on mouse brain and cardiac functions at baseline but accelerates the progression of neurobehavioral and cardiac impairments in AD mice, establishing for the first time that homeostatic activation of 26S proteasomes by p-S14-RPN6 or PKA protects against AD progression.

## METHODS

### Animals

The 5XFAD mouse in the C57BL/6J inbred background (RRID:MMRRC_034840-JAX) was originally obtained from The Jackson Laboratory. In the 5XFAD mice, the overexpression of both mutant human amyloid beta (A4) precursor protein 695 (APP) with the Swedish (K670N, M671L), Florida (I716V), and London (V717I) Familial Alzheimer’s Disease (FAD) mutations and human PS1 harboring two FAD mutations, M146L and L286V, is driven by the neural-specific elements of the mouse *Thy1* promoter ^44^. The creation and validation of the Rpn6^S14A^ knock-in (referred to as S14A hereafter) mice were recently reported ^41, 45^. The S14A mouse was created in and has been backcrossed into the C57BL/6J background for more than 10 generations. The 5XFAD mice and S14A mice were crossbred for necessary rounds to generate mice with appropriate genotypes for the experiments determining the effect of genetic blockade of p-S14-Rpn6 on the progression of AD reported here.

### Novel Object Recognition Test (NORT)

To test the different phases of learning and memory in mice we performed a novel object recognition test (NORT) as previously described^46, 47^. This test aimed to examine the recognition memory of mice by measuring the time spent exploring the novel object versus the familiar one. This test is based on the innate preference of the mice to explore the novel object. The object recognition protocol was adapted from the previously described. Mice were tested in a square wooden box (50 × 50 × 30 cm) located in a room with dim lighting. The objects used in our experiments are different in shape and texture: Lego towers (five bricks) and Falcon tissue culture flasks (25 cm2) filled with sand. The experiments were performed between 9 am to 6 pm. Mice were habituated to the empty box for 5 minutes the day before the test. On day 2, during the familiarization session, mice were presented with two similar objects in the box and permitted to explore freely for a maximum of 10 minutes. The objects were cleaned with 70% alcohol between trials to eliminate the olfactory cues. After an intersession interval (ISI) of 24 hours, one of the two objects was randomly replaced by a new one during the test session. Mice were placed back into the box and allowed to explore the two objects for a maximum of 10 minutes. Stop the experiment when there had been a 20-second exploration of both objects or when the maximum session time was reached in both familiar and test sessions. The amount of time spent exploring each object was recorded, and the relative exploration of the novel object was presented by a discrimination index (DI = (t^novel^ -t^familiar^) /(t^novel^ + t^familar^)). The criteria for exploration behavior were defined as whenever the mouse sniffed the object or touched the object while looking at it within a circle of 2 cm around the object. Climbing onto the object only does not count as exploration.

### Radial Arm Water Maze (RAWM) Tests

The 2-day RAWM test was used to evaluate spatial learning and memory. The protocol is followed as described previously^48, 49^. On the following day (day 2), each mouse was trained for 15 trials with the hidden platforms only and the experimenters were blinded to the genotypes of the mice at the time of testing^50^.

### Mouse Tissue Harvesting and Western Blot Analysis

The mice were euthanized once they reached the desired age, and the organs (heart and brain) were harvested. A portion of the frontal to mid-right brain cortex was subjected to isolation with RIPA (Radioimmunoprecipitation assay) buffer before sonication. The heart was cleaned and cut into two sections (upper and bottom). The upper section was subjected to protein analysis while the rest of the ventricles were stored for RNA analysis. The isolated protein (50-100μg) was resolved onto 12% SDS-PAGE gel electrophoresis (Bio-Rad, USA) and subsequently electroblotted onto either nitrocellulose or PVDF membrane. The membranes were blocked with 5% non-fat milk in TBS (Tris-Buffered Saline) supplemented with 0.1% Tween (TBST) at least for an hour at RT. The blots were probed with different primary antibodies and incubated overnight prior to adding horseradish peroxidase-conjugated secondary antibodies (rabbit/mouse) for an hour at room temperature. Anti-mouse β-actin was used as a loading control for the protein. The signals were detected using ECL kit (Thermo Fisher Scientific) under the ChemiDoc Imaging System (Bio-Rad, USA).

Western blot analysis for S14-phosphorylated RPN6 using a custom-made antibody as previously described ^42^.

### Proteasome Peptidase Activity Assays

The protocol for proteasome peptidase activity assays was described previously ^51^.

### Echocardiography

The procedure of serial echocardiography on mice mentioned in **Table 1** and **Table 2** was performed as previously described ^41, 51^.

**Table 1.** Number of WT and S14A/A mice subjected to echocardiography during aging.

|  |  |  |  |  |
| --- | --- | --- | --- | --- |
| **Age** | **WT** |  | **S14A/A** |  |
|  | male | female | male | female |
| 6 months | 10 | 11 | 5 | 11 |
| 12 months | 10 | 12 | 5 | 17 |
| 18 months | 10 | 8 | 5 | 14 |

**Table 2.** Number of mice subjected to echocardiography at each time point.

|  |  |  |  |  |  |  |  |  |  |  |
| --- | --- | --- | --- | --- | --- | --- | --- | --- | --- | --- |
|  | 1 month |  | 2 months |  | 4 months |  | 8 months |  | 12 months |  |
|  | m | f | m | f | m | f | m | f | m | f |
| WT | 10 | 3 | 12 | 8 | 11 | 11 | 11 | 11 | 11 | 11 |
| 5XFAD | 12 | 16 | 15 | 14 | 19 | 19 | 19 | 19 | 18 | 18 |
| 5XFAD::S14A/+ | 3 | 4 | 4 | 4 | 4 | 4 | 4 | 4 | 4 | 4 |
| 5XFAD::S14A/A | 9 | 4 | 6 | 3 | 7 | 4 | 7 | 4 | 7 | 4 |

M, male; f, female

### Statistical analysis

GraphPad Prism Statistical Software version 10.3.1 were used for statistical analysis and graphical display. All numerical data were presented as mean ± SEM or mean ± SD. Statistical methods used for each experiment are described in the figure legend. For the RAWM test, the average errors to find the platform were calculated in blocks of consecutive 3 trials. A two-way ANOVA followed by Tukey test was used for comparisons between different genotypes at each block. A repeated measure one-way ANOVA followed by Dunnett test was used for behavioral comparisons between different time points (blocks) within each genotype to test for learning/memory over time in that genotype. *P* value or adjusted *P* value <0.05 is considered significant.

## RESULTS

### 1. Characterization of Rpn6^S14A^ (S14A) mice at baseline

#### 1.1 Reduced proteasome peptidase activities in the cortex of S14A mice

First, we assessed the Ser14-phosphorylated Rpn6 protein levels in the cerebral cortex of S14A mice using Western blot analyses. As expected, Ser14-phosphorylated Rpn6 was undetectable in the homozygous S14A mice (S14A/A) and decreased by approximately 50% in the heterozygous S14A mice (S14A/WT), compared with the WT littermate controls (**Figure 1A)**. Then, we measured proteasome peptidase activities in the crude protein extracts from the brain cortex of WT and S14A/WT mice. Different from what we previously found in myocardium ^41^, all three peptidase activities in the cortex were significantly lower in the S14A/WT mice than in WT controls (**Figure 1B-1D**).

**Figure 1.**
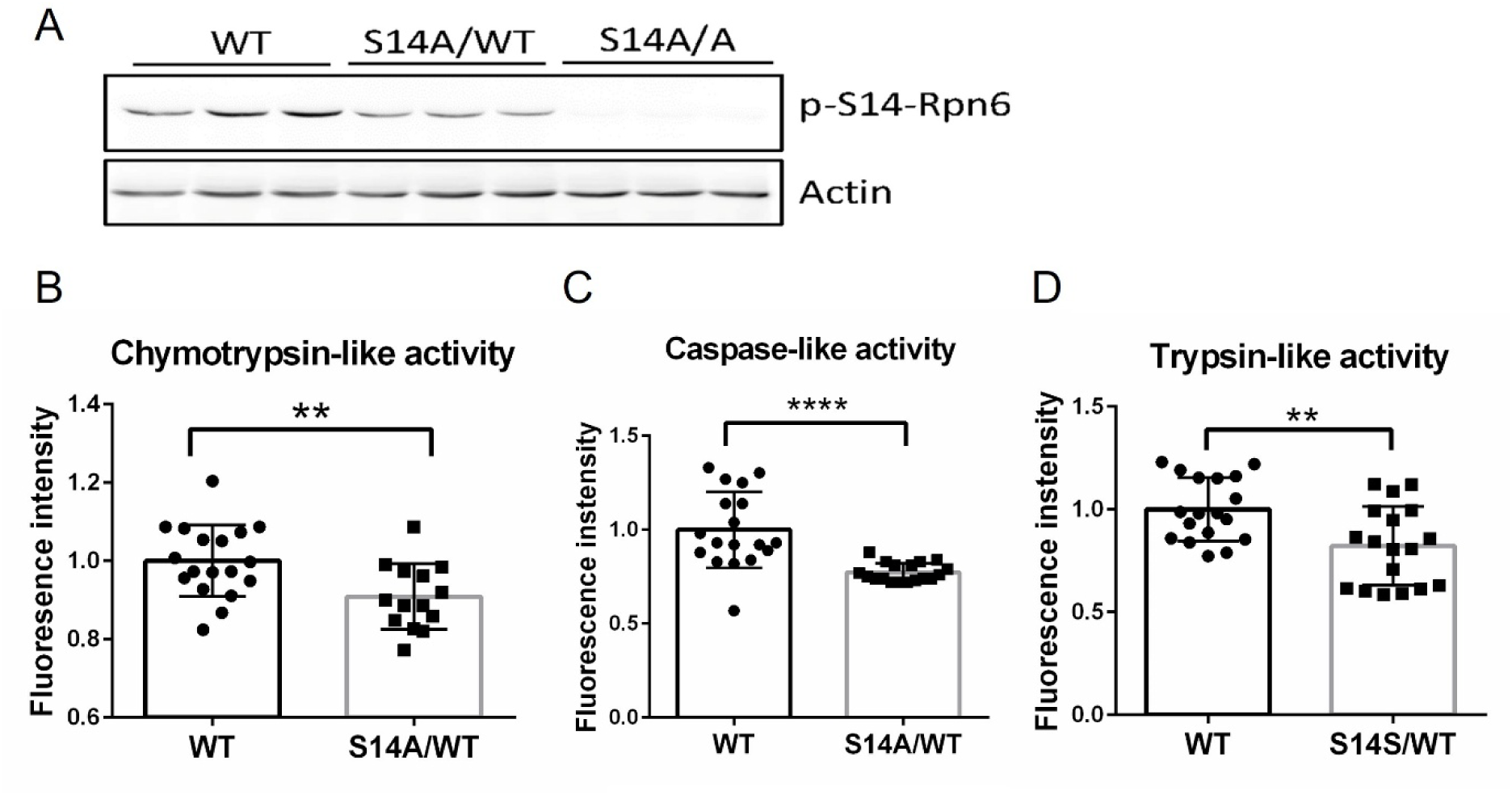
Changes of Ser14-phosphorylated Rpn6 (p-S14-Rpn6) protein levels and proteasome peptidase activities in the brain cortex of WT and S14A mice. **A**, Western blot analysis for p-S14-Rpn6 in the brain cortex of wild type (WT), heterozygous S14A (S14A/WT), and homozygous S14A (S14A/S14A) mice. Actin was immunoblotted as a loading control. **B**-**D**, Proteasome peptidase activity assays using crude protein extracts from WT and S14A/WT mice. Data are shown as mean ± SD; n = 6 for each group of mice. **p < 0.01, ***p < 0.001, unpaired Student’s *t*-tests.

#### 1.2 S14A mice showed a slight reduction in spatial learning and memory

Spatial learning and memory were assessed using the radial arm water maze (RAWM) and the results showed that the heterozygous S14A mice (S14A/WT) at 4 months showed a slight reduction in learning and memory at a later stage of the test, compared with the WT group (**Figure 2A**, Block 9). However, the time taken to reach the platform did not differ significantly between the two groups of mice (**Figure 2B**). For the homozygous S14A (S14A/S14A) mice, the pattern of error was similar to the S14A/WT; throughout the tests, the average error rate of the homozygous S14A mice appeared to be greater than that of the WT mice but the difference was close to reaching (p=0.06) but did not reach a statistical significance at any trials (**Figure 2C**). No difference of time used to find the platform between the S14A/S14A and the WT groups (**Figure 2D**).

**Figure 2.**
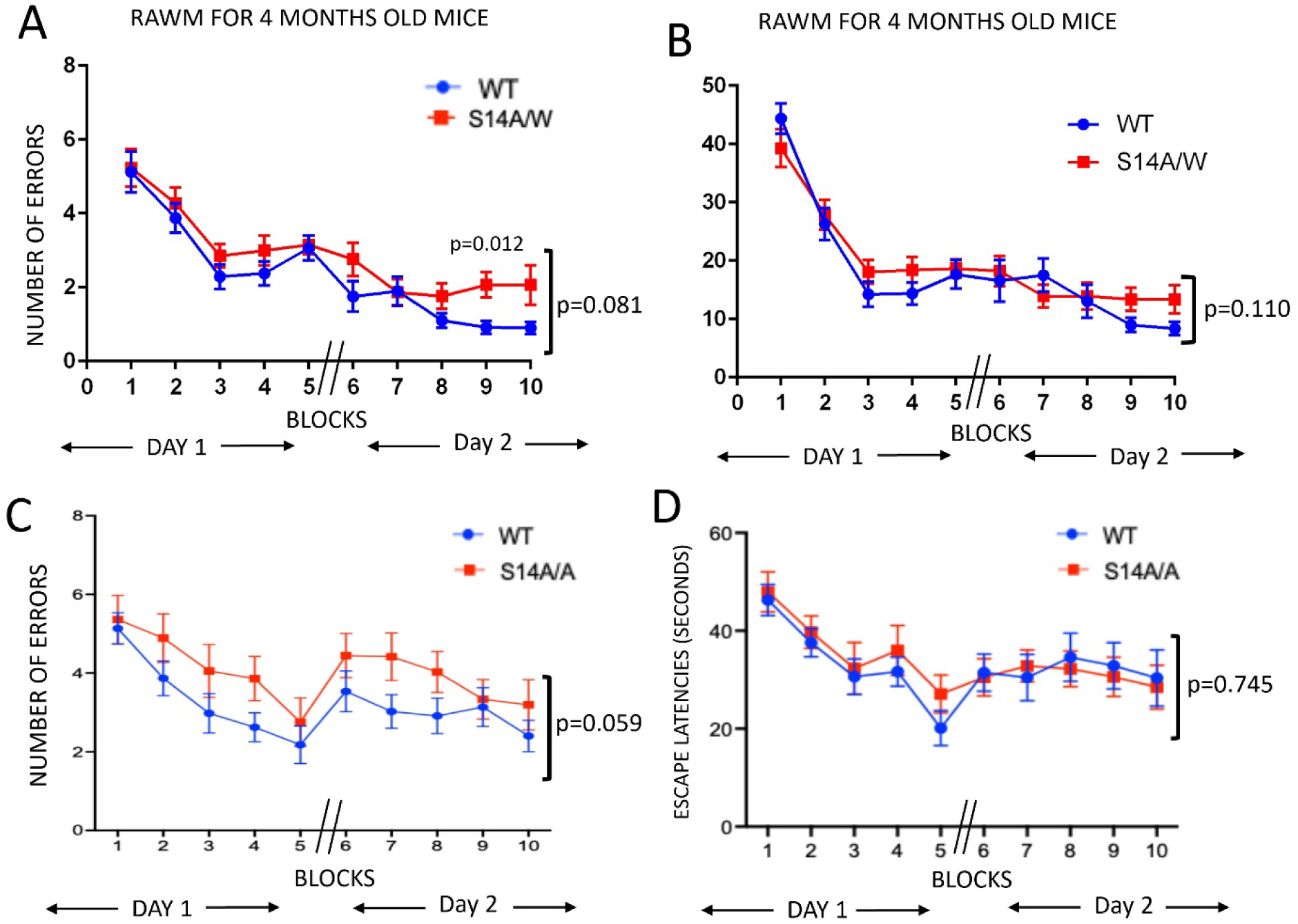
The radial arm water maze (RAWM) test results of mixed sex 4-month-old heterozygous S14A (S14A/+) mice (**A**, **B**; n=22) and homozygous S14A (S14A/A) mice (**C**, **D**; n=12) in comparison to wild type (WT) mice (n=16). Data are shown as mean±SEM. *P* values shown besides the bracket are for the genotype factor derived from two-way ANOVA.

### 2. Blockage of p-S14-Rpn6 exacerbates AD-caused learning and memory deficits

To decipher the role of p-S14-RPN6 in the progression of AD, we first examined cerebral cortex expression of Ser14-phosporylated Rpn6 in 5XFAD mice. Our results reveal a modest but statistically significant increase of Ser14-phosporylated Rpn6 in the AD mice compared with the WT controls (**Figure 3A, B**).

**Figure 3.**
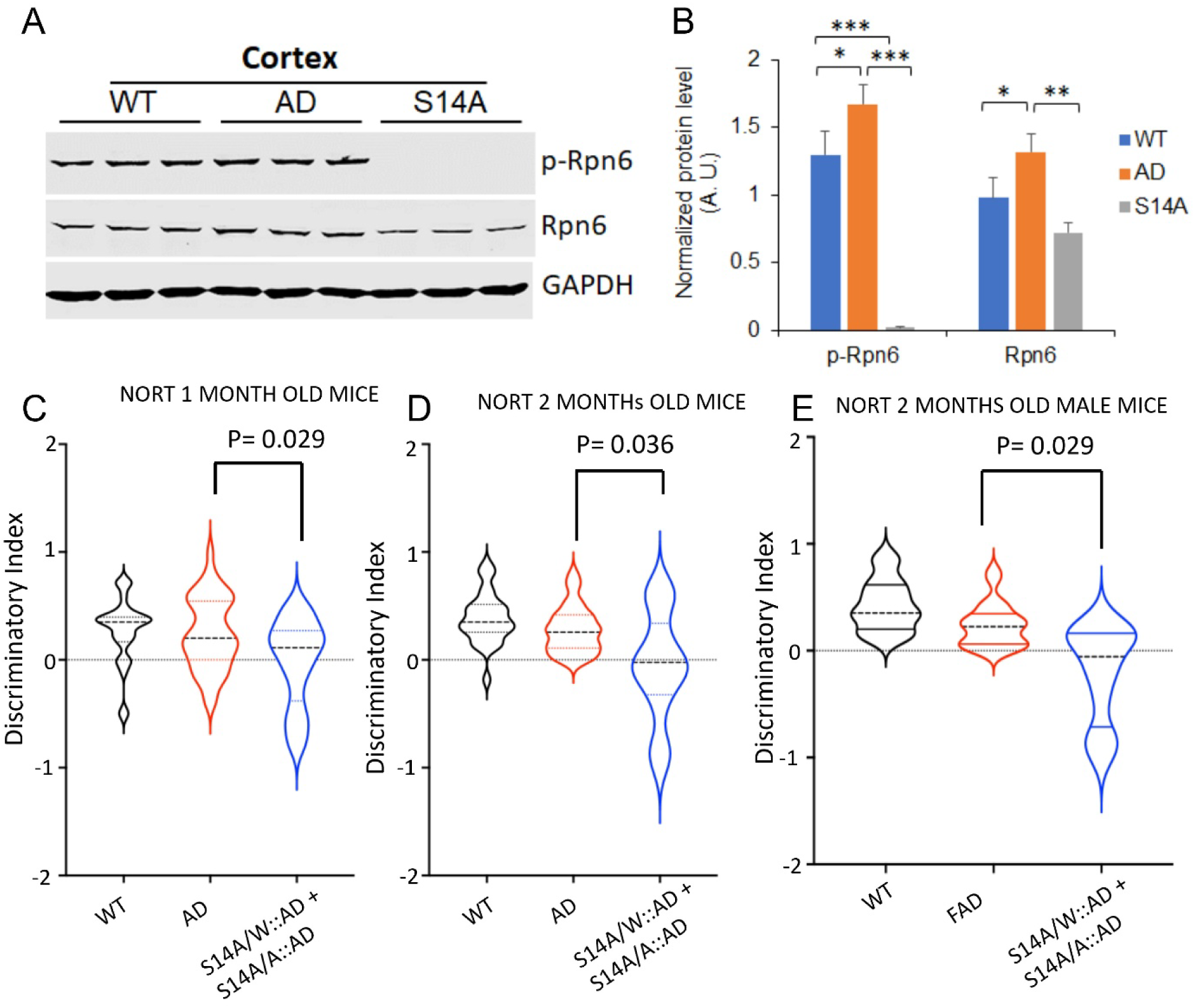
Ser14-phosphorylated Rpn6 is increased in the cerebral cortex of 5XFAD mice, and the novel object recognition test (NORT) reveals genetic blockade of the Ser14 phosphorylation of Rpn6 accelerates learning and memory deficits in 5XFAD mice. **A** and **B**, Representative images (A) and pooled densitometry data (B) of Western blot analysis for Ser14-phosphorylated Rpn6 (p-Rpn6) and total Rpn6 levels in the cerebral cortex of 6-week-old mice with the indicated genotypes. Data are shown as mean±SD; n=3 mice/group. *p<0.05, **p<0.01, ***p<0.001; one way ANOVA followed by Dunnett’s tests. **C**∼**E**, NORT showed that combined genotypes (n=22) of S14A/A::5XFAD and S14A/W::5XFAD (referred to as S14AA/AW::AD in this figure) exhibited significant short-time memory deficits in comparison to 5XFAD (n=43) and WT (n=29) mice at 1-month (**C**) to 2-month (**D**) timepoints. Data are shown as mean ± SEM. *p<0.05. (**E)** The results show that the cognition deficit in *male* mice is highly significant (p<0.001) at 2 months duration. The significance was calculated using one-way ANOVA followed by Dunnett’s test in GraphPad Prism.

To test the role of Ser14-phosporylated Rpn6 in AD, we crossbred S14A into the 5FXFAD mice and performed the novel object recognition test (NORT) on the resultant littermate mice. NORT is known to assess learning and short memory. We found that combined genotypes (n=22) of S14A/A::5XFAD and S14A/W::5XFAD (referred to as S14AA/AW::AD in **Figure 3C-3E**) exhibited significantly lower discriminatory index in comparison to the 5XFAD mice at 1 month, the youngest age examined (**Figure 3C**) and 2 months of age (**Figure 3D**). Interestingly, the *males* but not females of the combined genotypes (S14A/S14A::AD and S14A/WT::AD mice) revealed a more pronounced reduction of the discriminatory index at 2 months compared to AD (**Figure 3E**). These NORT results indicate that genetic blockage of p-S14-Rpn6 accelerates AD-caused learning and memory deficits.

Next, we further measured the spatial learning and memory in the heterozygous S14A-coupled 5XFAD (S14A/W::AD) mice and the 5XFAD mice at 2 months of age using the radial arm water maze (RAWM) test. Our results showed that the S14A/W::AD mice made dramatically more errors at the later stage of the test (block 5 during day 1 and every block during day 2), compared to the 5XFAD mice (**Figure 4A**), although the time spent finding the platform did not show any significant difference between the two groups (**Figure 4B**). This is much earlier than the 5XFAD mice that show cognitive impairments at 3-6 months of age, suggesting that blockage of p-S14-Rpn6 exacerbates AD-caused learning and memory deficits.

**Figure 4.**
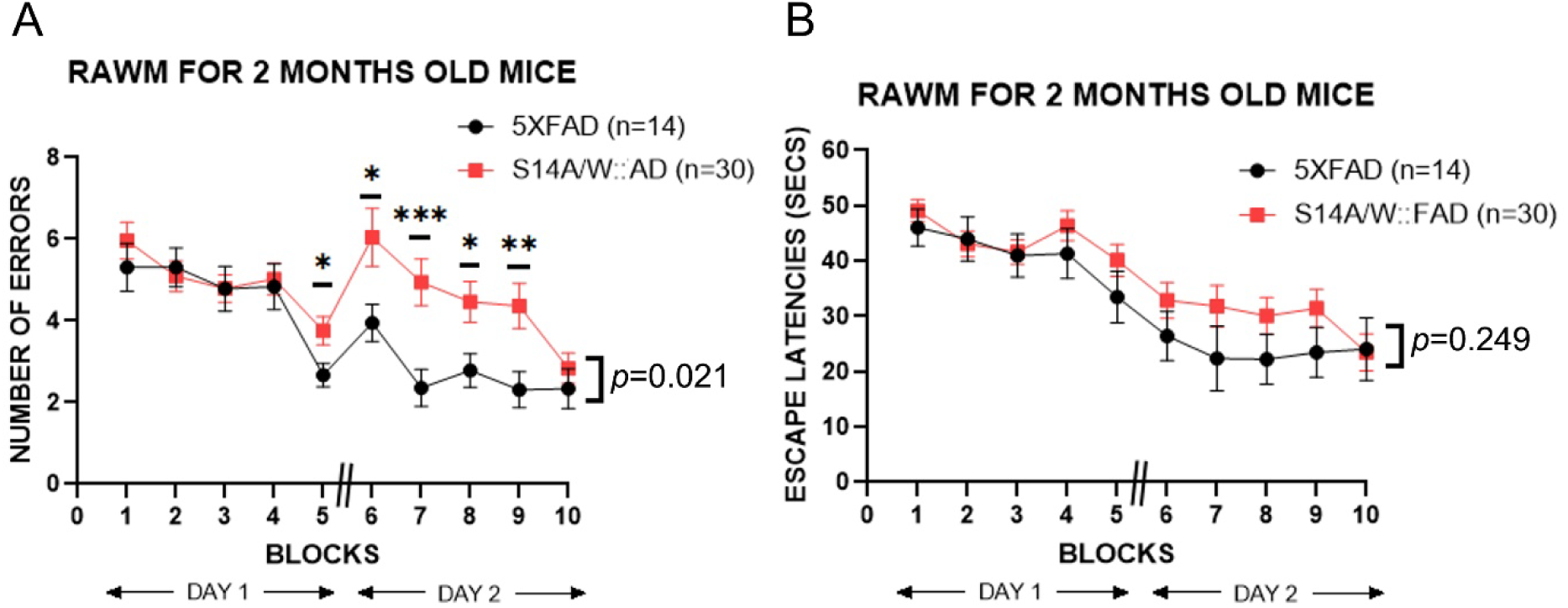
The radial arm water maze (RAWM) test results from the comparison between the heterozygous S14A coupled 5XFAD (S14A/W::AD) mice (n=30 mice) and the 5XFAD mice (n=14 mice) at 2 months of age. Mice of both sex were used. **A**, the number of errors made to find the platform. **B**, the escape latency (time spent finding the platform). Mean±SEM at each block are presented. Each block is an average of 3 trials. * p<0.05, **p<0.01, and ***p<0.001. **//** means a 24-hour break. Significant and noteworthy *P* values are shown above the block are derived from two-way ANOVA followed by Tukey’s tests; the *P* value shown to the right side of the bracket by Block 10 is from the genotype factor of the two-way ANOVA. The same conventions apply to Figures 5∼**7**.

At 4 months, the learning and memory deficits in the S14A/WT::AD (**Figure 5A**, **5B**) and S14A/A::AD mice (**Figure 6A**, **6B**) remained but appeared less prominent compared to 2 months. This might be caused by repeated training and tests within a relatively short period to the same cohort of mice. It is noted that the behavioral deficits in *male* mice are also prominent at the age of 4 months for both S14A/WT::AD (**Fig. 6A**) and S14A/A::AD mice (**Fig. 6B**).

**Figure 5.**
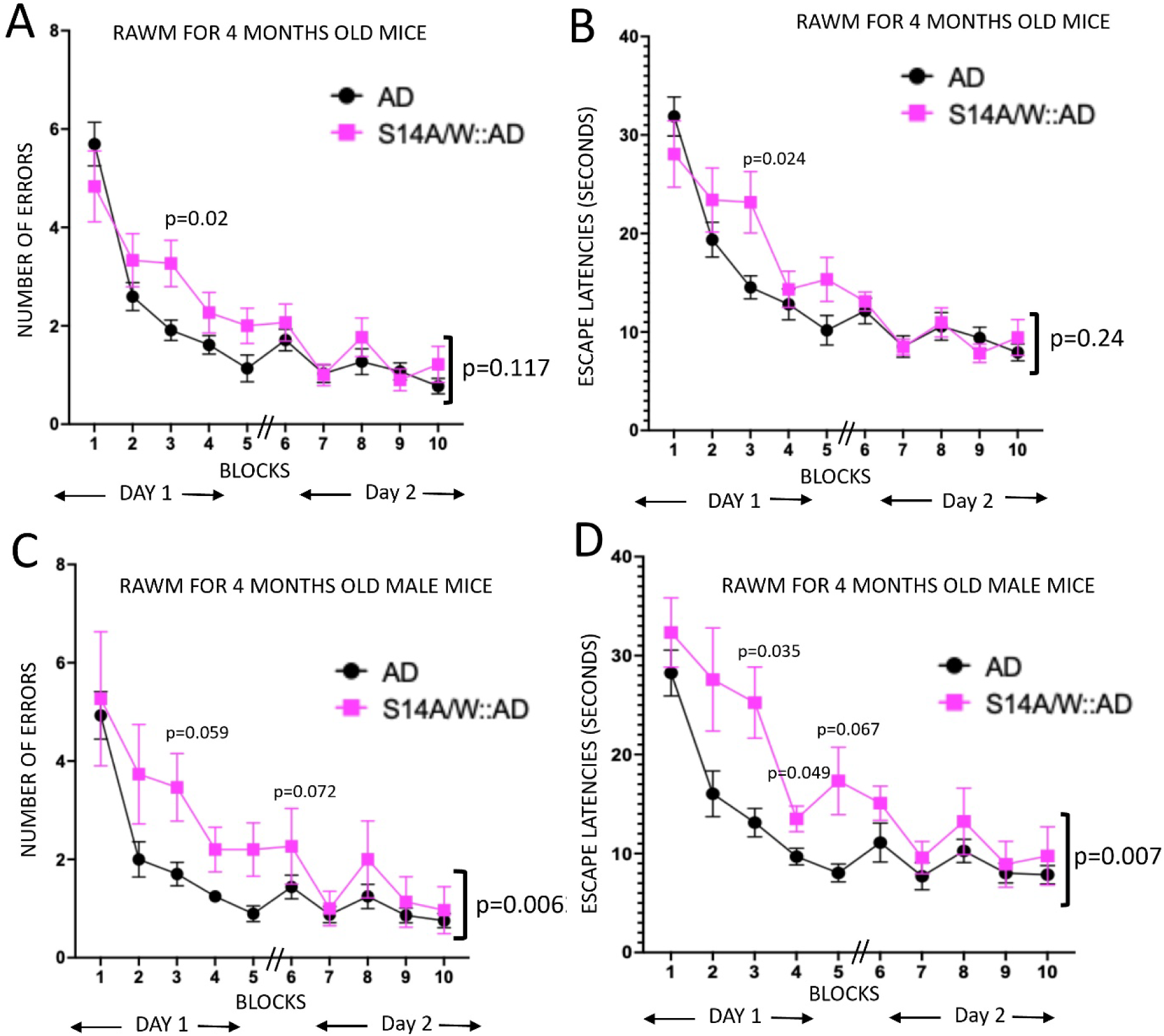
The performance differences in the radial arm water maze (RAWM) tests between the 5XFAD mice (5XFAD) and the heterozygous S14A coupled with 5XFAD mice (S14A/W::AD) at 4 months of age. **A** and **B**, Number of errors (A) and escape latency (B) exhibited by male and female mice combined. N=37 mice for 5XFAD, n=12 mice for S14A/W::AD. **C** and **D**, Number of errors (C) and the escape latency (D) exhibited by only male 5XFAD (n=17) and S14A/W::AD (n=5) mice. Mean±SEM. Each block is an average of 3 trials. * p<0.05, **p<0.01, and ***p<0.001. ns means not significant between groups while Sig. means significant. The interval between Block 5 and 6 (**//)** was 24 hours. *P* values shown above the block are derived from two-way ANOVA followed by Tukey’s tests. The *P* value shown to the right side of the bracket by Block 10 is from the genotype factor of the two-way ANOVA.

**Figure 6.**
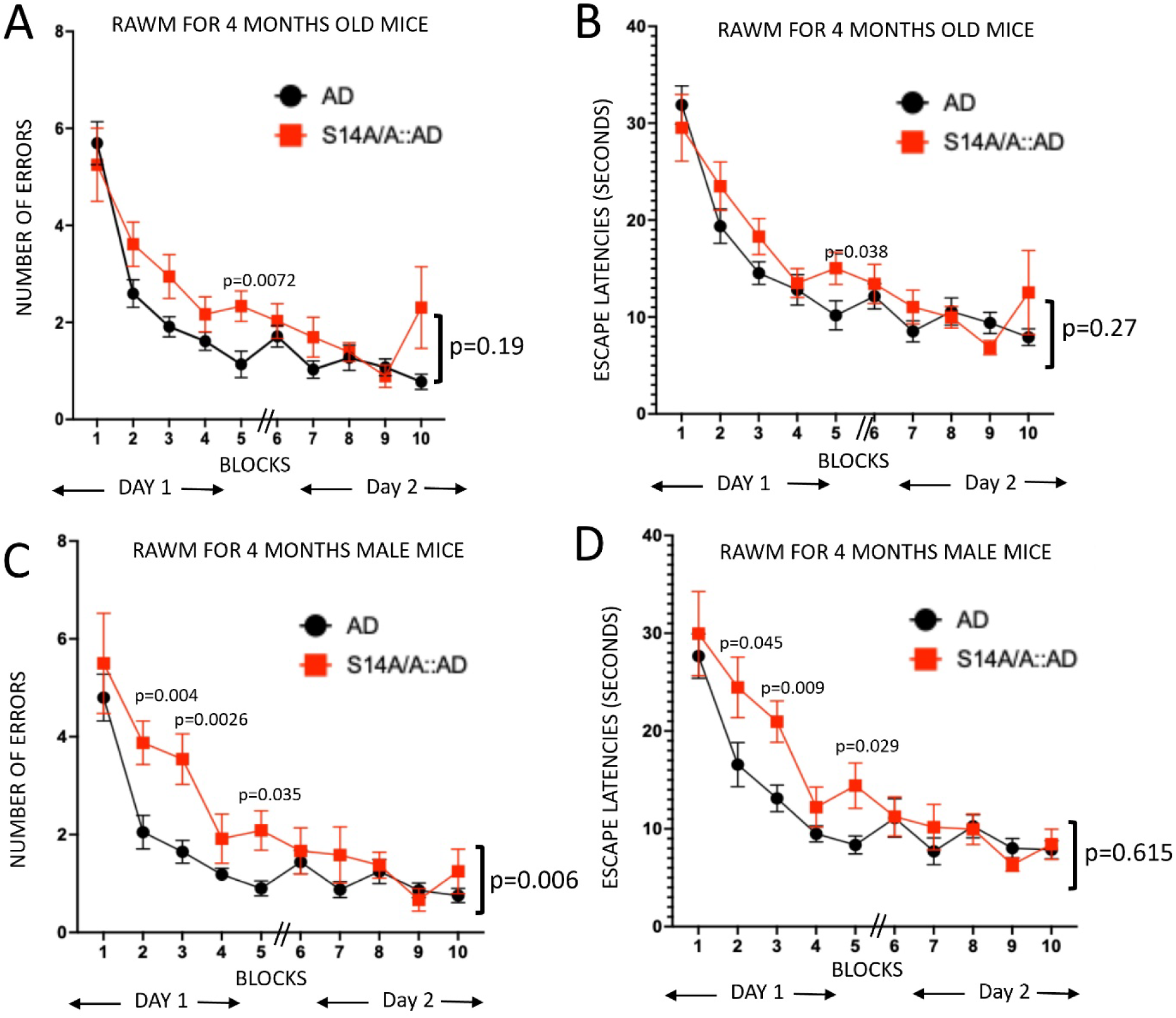
The performance differences in the radial arm water maze (RAWM) tests between the 5XFAD mice (5XFAD) and the homozygous S14A coupled with 5XFAD mice (S14A/A::AD) at 4 months of age. **A** and **B**, Number of errors (A) and escape latency (B) exhibited by male and female mice combined. N=37 mice for 5XFAD, n=10 mice for S14A/A::AD. **C** and **D**, Number of errors (C) and escape latency (D) exhibited by male 5XFAD (n=19) and S14A/A::AD (n=7). Mean±SEM. Each block is an average of 3 trials. * p<0.05, **p<0.01, and ***p<0.001. ns means not significant between groups while Sig. means significant. The interval between Block 5 and 6 (**//)** was 24 hours. *P* values shown above the block are derived from two-way ANOVA followed by Tukey’s tests. The *P* value shown to the right side of the bracket by Block 10 is from the genotype factor of the two-way ANOVA.

The error values gathered by RAWM assessment in 1-year-old S14A/A::AD mice (**Fig. 7A**, left) and S14A/W::AD **(Fig. 7A**, right) are higher than AD mice. The similar results were also observed in the male S14A/A::AD mice (**Fig. 7B**, left) and S14A/W::AD **(Fig. 7B**, right) mice. On day one (block 1-5; **Fig7A**), some of the year-old mice S14A/A::5xFAD mice failed to locate the platform during the protocol and hence caused variation in errors. However, most of the rodents (male and female) of this genotype located the hidden platform within the timeline after making substantial errors. Although, it is conspicuous that S14A/W and S14A/A crossed with 5xFAD mice exhibited behavioral deficits during 4 months RAWM trials but none of the participants of the cohort suffered from seizures. During our observation in year old mice we have noted a male (S14A/A::5xFAD) and three females (one S14A/W::5xFAD and two S14A/A::5xFAD) mice suffered from seizures during and after RAWM trails. Exhibition of seizures is a target component of AD progression as seen in 22% of AD patients, also aging APP/PS1 mice have been documented to suffer from seizures^52, 53^. Thus, blockage of phosphorylation of PSMD11 manifests in severe cognitive deficits akin to AD.

**Figure 7.**
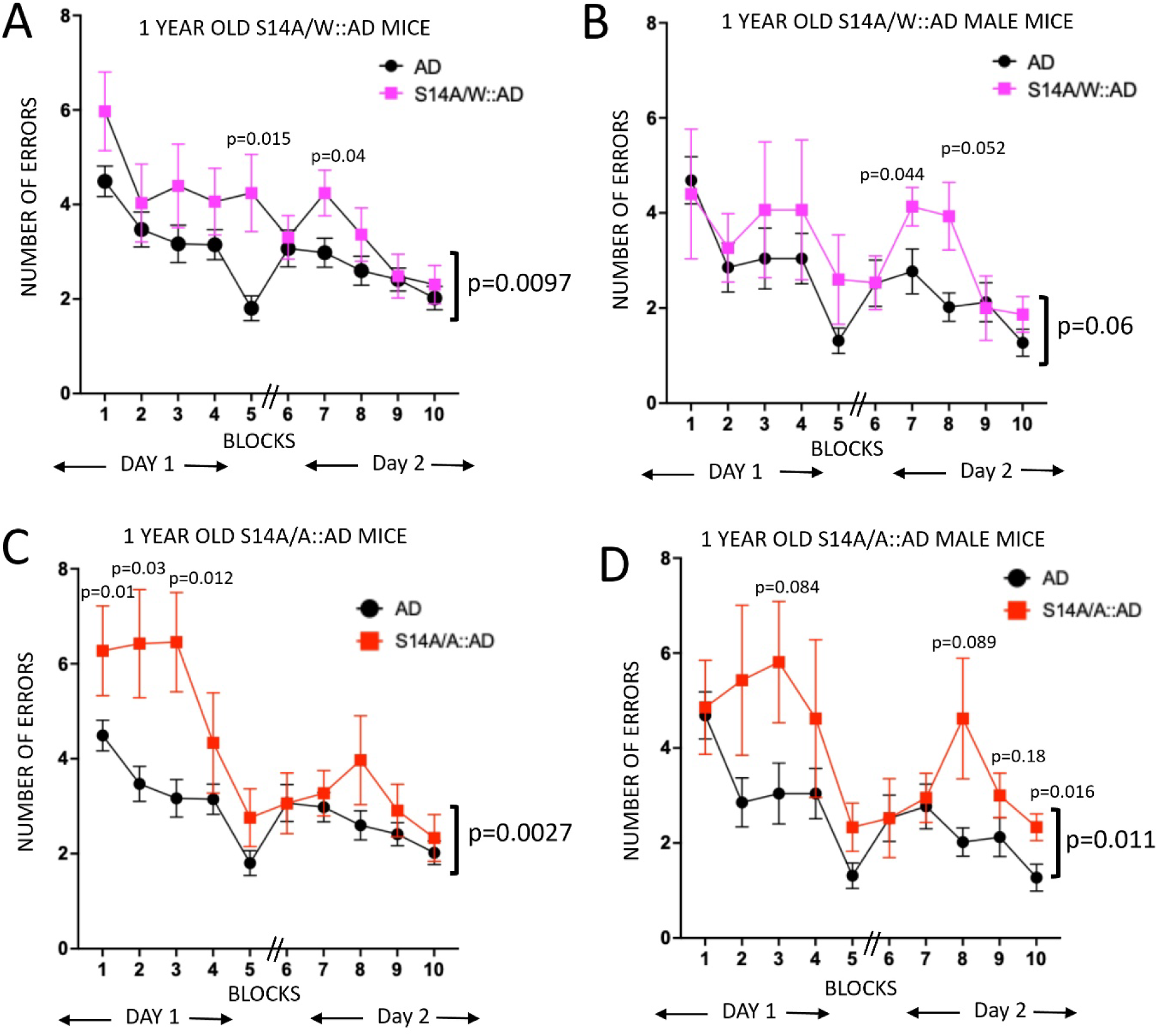
Comparison of the numbers of errors from RAWM tests between the 5XFAD mice (AD) and the heterozygous S14A coupled 5XFAD mice (S14A/W::AD, **A** and **B**) or homozygous S14A coupled 5XFAD mice (S14A/A::AD, **C** and **D**) at 12 months of age. **A** and **C**, males and females combined; n=11 for S14/W::AD, n=11 for S14A/A::AD, n=34 for AD. **B** and **D**, males only; n=5 males for the S14A/W::AD group, n=7 males for the S14A/A::AD group, and n=15 males for the AD group. Mean±SEM. Each block is an average of 3 trials. * p<0.05, **p<0.01, and ***p<0.001. ns means not significant between groups while Sig. means significant. The interval between Block 5 and 6 (**//)** was 24 hours. *P* values shown above the block are derived from two-way ANOVA followed by Tukey’s tests. The *P* value shown to the right side of the bracket by Block 10 is from the genotype factor of the two-way ANOVA.

### 3. Genetic blockade of RPN6-Ser14 phosphorylation causes modest cardiac malfunction in mice during ageing

Cardiac morphometry and function of the homozygous S14A mice at the age of 1 through 7 months had been characterized via serial Echo and no remarkable abnormalities were detected ^41, 45^. To determine whether genetical blockade of p-S14-RPN6 alters mouse cardiac morphometry and functions at a later age, here we have further performed serial Echo on homozygous S14A (S14A/A) and WT control mice at 6, 12, and 18 months of age. Consistent with our prior reports, our analyses of the echocardiographic data reveal that S14A mice do not display significant differences in cardiac functional and morphometric parameters until 12 months of age, compared with the WT control group (**Figure 11**). The size of the heart (LV chamber diameters and volumes, wall thickness, LV mass) and cardiac output (stroke volume, output per minute) are known to be proportional to the body weight (BW); thus, we normalized the related parameters to BW to minimize the potential impact of BW. There were no statistically significant differences in any of the morphometric and functional parameters between the WT and S14A/A groups at 6 months. By 12 months, only the BW-adjusted LV end-diastolic diameter (LVID;d; **Figure 11H**) and LV end-diastolic volume (EDV; **Figure 11C**) show a slight but statistically significant reduction in the S14A/A mice compared with the WT control group. By 18 months, a modest but statistically significant decline of cardiac function as reflected by reduction of SV/BW and cardiac index (CO/BW) was observed in the S14A/A mice compared with the WT group (**Figure 11E** and **11F**). This modest cardia impairment does not appear to be caused by systolic malfunction because the ejection fraction (EF; **Figure 11D**) and fractional shortening (data not shown) did not show a significance change, whereas a mild diastolic malfunction might be blamed as the LVID;d/BW and EDV/BW tend to smaller in the S14A/A mice than in WT mice throughout the study duration and the differences achieved statistical significance at 12 months (**Figure 11H** and **11C**). Notably, throughout the duration of the study, no statistically significant differences in LV posterior wall thickness at the end of diastole (LVPW;d/BW; **Figure 11G)** or estimated LV mass (LV mass/BW; **Figure 11I**) were detected between S14A/A and WT mice, indicating that genetic blockade of p-S14-Rpn6 does not cause cardia hypertrophy in mice.

### 4. The 5XFAD mouse heart displays early-onset increases in Aβ and decreases in proteasome activities and late-onset cardiac malfunction

Cardiac malfunction is often associated with AD and amyloid beta (Aβ) protein aggregates were found to be present in the hearts of AD patients with a primary diagnosis of AD and impair cardiomyocyte functions ^54^, suggesting a brain-to-heart link in AD pathogenesis. Moreover, a recently reported study detected remarkable declines in cardiac systolic function in the 5XFAD mice as early as 2 months of age^55^. Therefore, we examined the protein levels of the amyloid precursor protein (APP) and Aβ in both the brain cortex and myocardium of the 5XFAD mice at 4-week of age. As expected, an increased expression of APP proteins was detected only in the brain not in myocardium, compared with the WT controls (**Figure 9A∼9D**). However, an unexpected, marked increase in Aβ was detected in the heart but not the brain of 5XFAD mice at this age (**Figure 9A**, **9B**, and **9E**). Consistent with increased intracellular Aβ impairs the proteasome, myocardial proteasome peptidase activities were significantly decreased in the 5XFAD mice (**Figure 9F**). This prompted us to perform a thorough echocardiographic study on these mice as we determine the impact of genetic blockade of -S14-RPN6 on AD progression (see next Section).

We performed serial Echo on a cohort of 5XFAD mice and their WT littermates. Starting between 4 and 8 months of age, the 5XFAD mice showed decreased body weight (BW). Thus, for LV wall thickness and chamber size as well as SV and CO assessments, we used BW-adjusted parameters for comparisons. Analyzing the echocardiograms collected at 1, 2, 4, 8, and 12 months of age, we found that significant increases in BW-adjusted LVPW;d (**Figure 10B**), LVPW;s (**Figure 10C**), LVID;d (**Figure 11A**), and LVID;s (**Figure 11B**), signs of cardiac hypertrophy, were evident in the 5XFAD mice at 12 months, compared with the age-matched WT control group. Compared with the WT group, the AD mice did not show discernible functional deficits until 8 months of age when their cardiac index (CO/BW) were significantly lower (**Figures 11I**, **13D**, **14D**). Interestingly, no statistically significant differences in fractional shortening (FS) and ejection fraction (EF), two commonly used indies for systolic function, were detected between the AD and WT groups throughout the 12 months of age (**Figures 11C**, **11F, and Panels A and B of Figures 12**∼**14**), suggesting that diastolic impairment may account for the decreased cardiac index discerned at 12 months.

### 5. Genetic blockade of RPN6-Ser14 phosphorylation exacerbates cardiac malfunction in AD mice

To establish the role of p-S14-RPN6 in AD-associated cardiac pathogenesis, we performed serial echocardiography on the same cohorts of mice as used for the neurobehavior tests (**Figures 2**∼**7**), that were derived from introduction of the S14A allele into the 5XFAD mice through multiple rounds of crossbreeding. Since our earlier studies have revealed that homozygous S14A mice do not display significant changes in the cardiac index (CO/BW) during the first 12 months of age (**Figure 8F**), we did not include the mouse groups with heterozygous S14A (S14A/W) or homozygous S14A (S14A/A) alone in this experiment to reduce the cost. Thus, the main comparisons of this segment of the study focus on heterozygous S14A coupled 5XFAD (S14A/W::AD) and homozygous S14A coupled 5XFAD (S14A/A::AD) in comparison with WT and the 5XFAD groups (**Figures 10**∼**14**).

**Figure 8.**
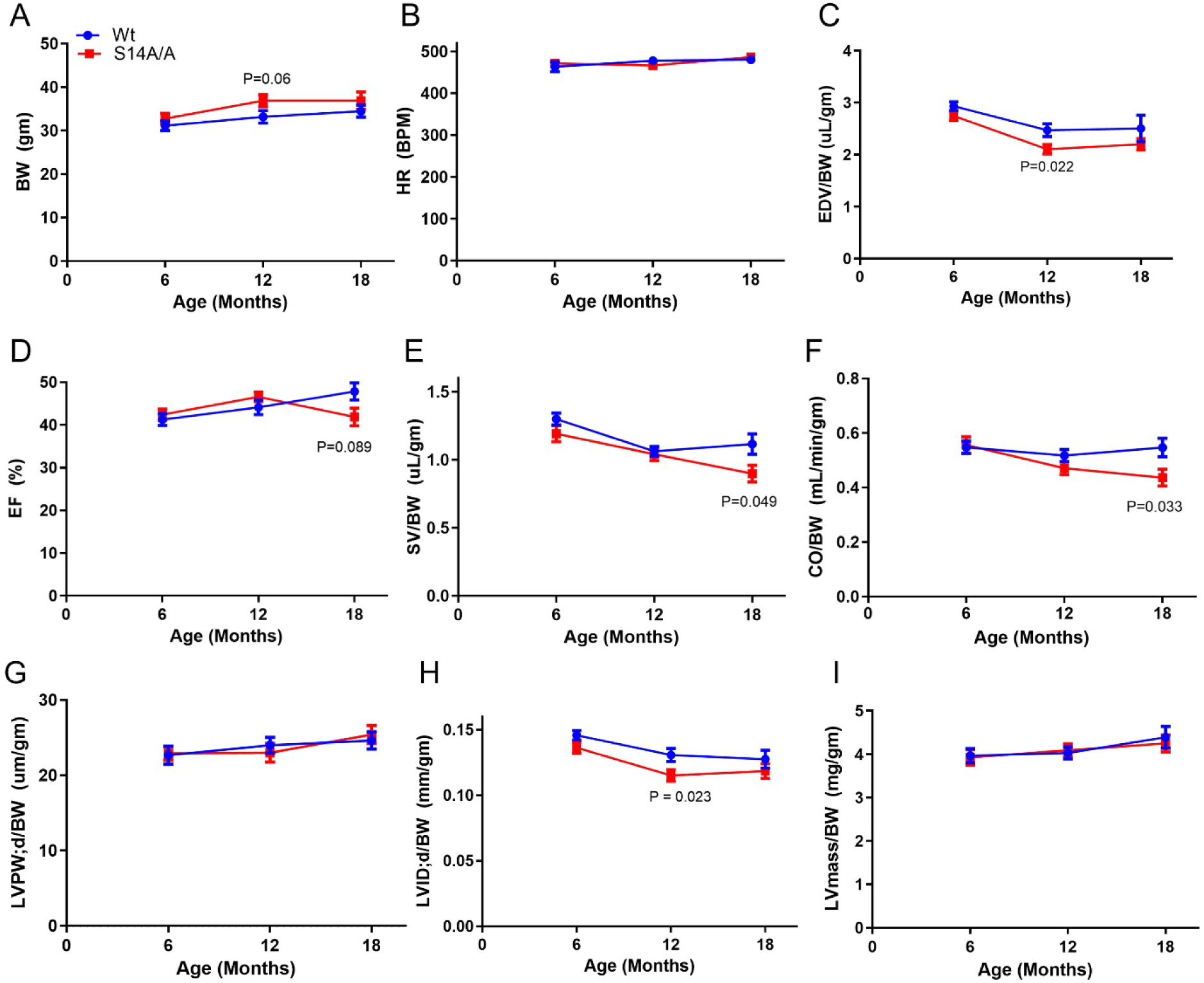
Serial echocardiographic characterization of the effects of genetical blockade of Ser14 phosphorylation of Rpn6 on mouse cardiac morphology and functions. The 2D image guided M-mode echocardiography was performed on homozygous S14A (S14A/A) and wild type (Wt) control mice at 6, 12, and 18 months of age. Body weight (BW) was measured immediately before echocardiography. HR, heart rate, EDV, left ventricular (LV) end-diastolic volume, EF, ejection fraction, SV, stroke volume; CO, cardiac output per minute; LVPW;d, LV posterior wall thickness at the end of diastole; LVID;d, LV end-diastolic diameter; LV mass, estimated LV mass. Mean±SEM; refer to Table 1 for the number of male and female mice included. *P* values shown were obtained from multiple *t*-tests.

Neither S14A/W nor S14A/A discernibly altered the reduction of body weight or the changes of LVPW;d and LV mass in 5XFAD mice (**Figure 10A**, **10B**, **10D**). As mentioned earlier, 5XFAD mice do not show discernible changes in EF and FS throughout the 12 months when compared with the WT, but both S14A/W::AD and S14A/A::AD mice, especially the latter, displayed significant reductions of FS, EF, SV/BW, and CO/BW as early as 4 months of age (**Figures 10**∼**14**). Notably, S14A/A::AD mice exhibited significantly lower FS, EF, SV/BW, and CO/BW than the littermate 5XFAD mice as early as 4 months (CO/BW) or 8 months of age (**Figures 10-14**). These results indicate that the partial or complete blockade of homeostatic pS14-Rpn6 accelerates and exacerbates cardiac malfunction in 5XFAD mice, thereby demonstrating that Ser14-phohsphorylation in RPN6 can protect against systolic malfunction during AD progression.

## DISCUSSION

AD is a multifactorial disease with multiple symptoms, exhibiting pathologies in both the brain and other organs including the heart^56^. However, the relationship of these peripheral pathologies to AD remains obscure. Augmentation of the cAMP/PKA pathway can activate the 26S proteasome via p-S14-RPN6 but the pathophysiological role of p-S14-RPN6 in AD has not been established. Here, we have not only identified a potential brain-to-heart axis in AD pathogenesis but also demonstrated for the first time that genetic blockade p-S14-Rpn6 accelerates the progression of both neurobehavioral deficits and cardiac malfunction in the 5XFAD mice, thereby establishing for the first time that homeostatic activation of 26S proteasomes by p-S14-RPN6 or by basal PKA activity protects against AD progression in both the brain and heart.

### New evidence for a brain-heart axis in AD pathogenesis

Although it has been frequently observed that cardiovascular diseases, such as decreased cerebral blood flow and heart failure, have a high incidence of coexisting with AD among older patients^57, 58^, the relationship between these peripheral symptoms/pathologies and AD remains unclear.^56, 59^ Most AD is associated with ageing which by itself affects most, if not all, organs, making it difficult to sort out the relationship between the pathogenesis of peripheral organs and the central nervous system (CSN) in AD development. Similarly, the genetic mutations-linked familial AD could potentially affect both the CSN and peripheral organs including the heart. For example, mutations in familial AD (FAD) related genes such as PSEN1 and PSEN2 are also linked to dilated cardiomyopathy and heart failure in humans.^60^ In the 5XFAD mouse model used in the present study, the expression of the five disease-causing transgenic mutations is driven by a the Thy1 promoter, which is brain-specific. Indeed, our results show that APP expressions for example are only increased in the brain not in the myocardium (**Figures 9A**, **9C**, and **9D**). However, Aβ (the derivative APP) in myocardium of the 5XFAD mice was multi-folded increased compared with the WT mice at 1 month of age. This increased Aβ in myocardium even occurred before the increases of Ab in the brain were discernible (**Figure 9B**, **9E**). This striking and unexpected findings provide new support that the Aβ originated form the brain can enrich in a peripheral organ such as the heart and thereby play a pathogenic role. Interestingly, contradicting to a prior report which shows an early onset cardiac systolic malfunction in a 5XFAD mouse model^55^, we found that a reduced cardiac index did not become discernible until 8 months of age and a deficit in systolic function was not detected throughout the entire 12 months of age (**Figure 11**), suggesting that the late-onset cardiac malfunction of the 5XFAD mice is likely caused by diastolic malfunction rather than systolic impairment. To support this proposition, <u>Troncone</u> *et al*. reported patients with AD displayed diastolic dysfunction^54^.

**Figure 9.**
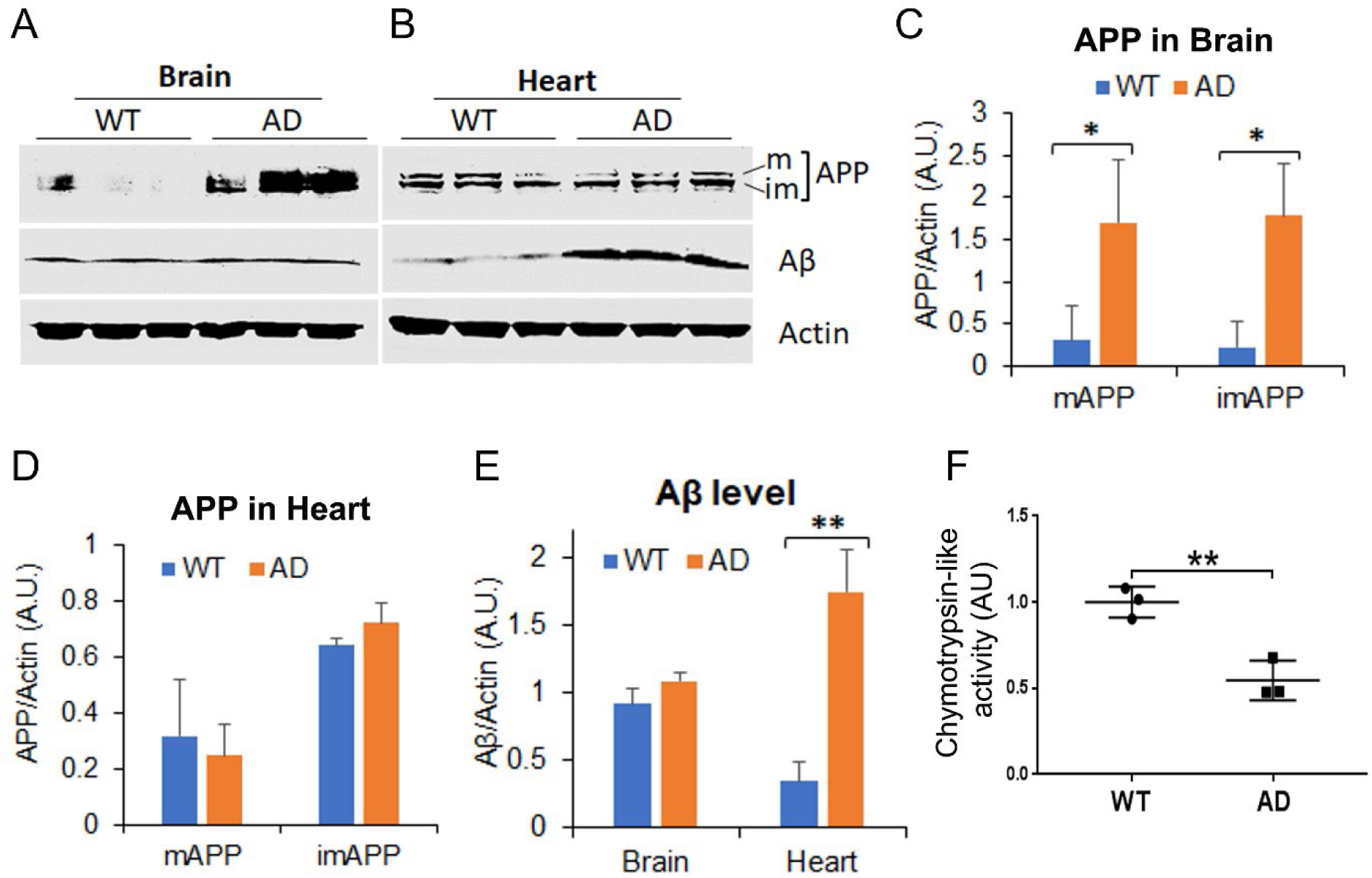
Increased Aβ and reduced proteasome peptidase activity in the myocardium of 5XFAD mice at 4 weeks of age. **A**∼**E**, Representative images (**A**, **B**) and pooled densitometry data (**C**∼**E**) of Western blot analyses for APP and Aβ levels in mixed-sex AD and WT brain cortex and ventricular myocardium. m, mature; im, immature. **F**, Proteasome chymotrypsin-like activity. Crude myocardial protein extracts from mixed-sex AD and WT mice and synthetic fluorogenic peptide substrate were used. Mean±SD; n=3 mice/group. \**p*<0.05, \*\**p*<0.01; Student’s *t*-test.

### Ser14-RPN6 phosphorylation protects against both neurobehavioral and cardiac malfunction during AD progression

The relatively late onset of cardiac malfunction (6 months of age) compared to the early onset increases of Aβ in myocardium (1 month of age) of the 5XFAD mice suggests the presence of powerful mechanisms that counter the pathogenic actions of Aβ. The present study demonstrates that homeostatic p-S14-RPN6 and associated proteasome activation are among the key mechanisms that protect against AD pathogenesis in both the brain and the heart. This claim is compellingly supported multiple lines of complementary evidence reported here. First, Ser14-phosphorylated Rpn6 was significantly increased in both myocardium (*data not shown*) and brain cortex (**Figure 3**) of 5XFAD mice; second, cerebral cortex proteasome activities were significantly decreased in S14A mice (**Figure 1**), although this did not appear to in the myocardium of S14A mice^41^; and third, genetic blockade of p-S14-Rpn6 aggravated learning and memory deficits (**Figures 2∼7**) and remarkably accelerates and exacerbates cardiac malfunction (**Figures 10∼14**) in the5XFAD mice. These findings are in full agreement with another related report where we have demonstrated that homeostatic activation of 26S proteasomes via p-S14-Rpn6 protects against both neural and cardiac pathology in a tauopathy mouse model (https://www.biorxiv.org/content/10.1101/2025.03.24.645024v1). Our findings are also consistent with the proposition that proteasome activation via p-S14-RPN6 may be a key mechanism underlying the protection against AD by cAMP/PKA augmentation although this remains to be formally tested.

**Figure 10.**
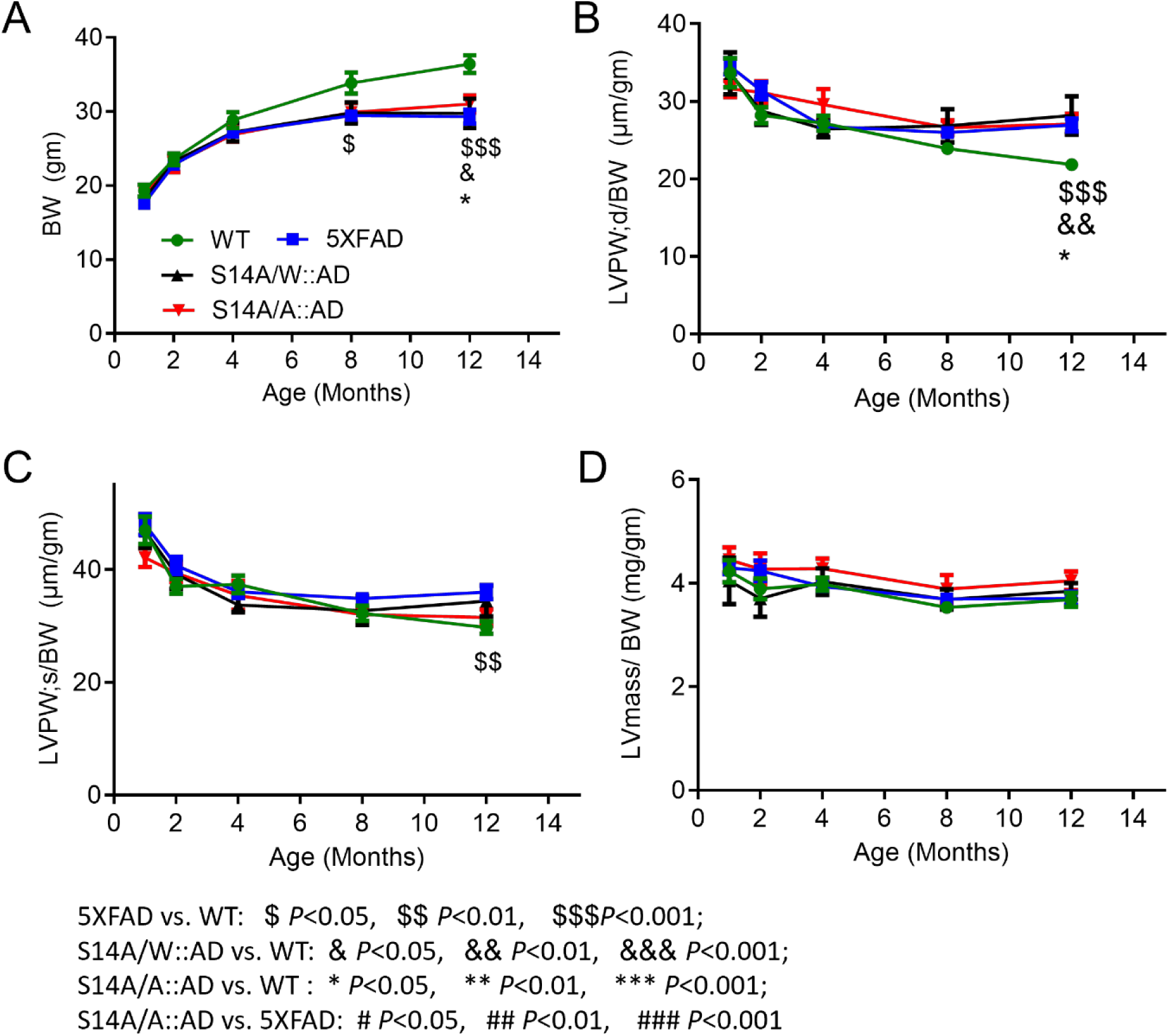
The time course of changes in body weight (BW), left ventricular (LV) posterior wall thickness at the end of diastole (LVPW;d) and at the end of systole (LVPW;s), and estimated LV mass (LVmass) in mice with the indicated genotypes as derived from serial echocardiography. Both males and females are included (see **Table 2** for sex distribution). Mean±SEM are plotted at each time point; WT, wild type; S14A/W::AD, heterozygous S14A coupled with 5XFAD; S14A/A::AD, homozygous S14A coupled with 5XFAD. The conventions denoting the genotypes/groups and symbols denoting pair-wise comparison *P* values also apply to Figure 11.

**Figure 11.**
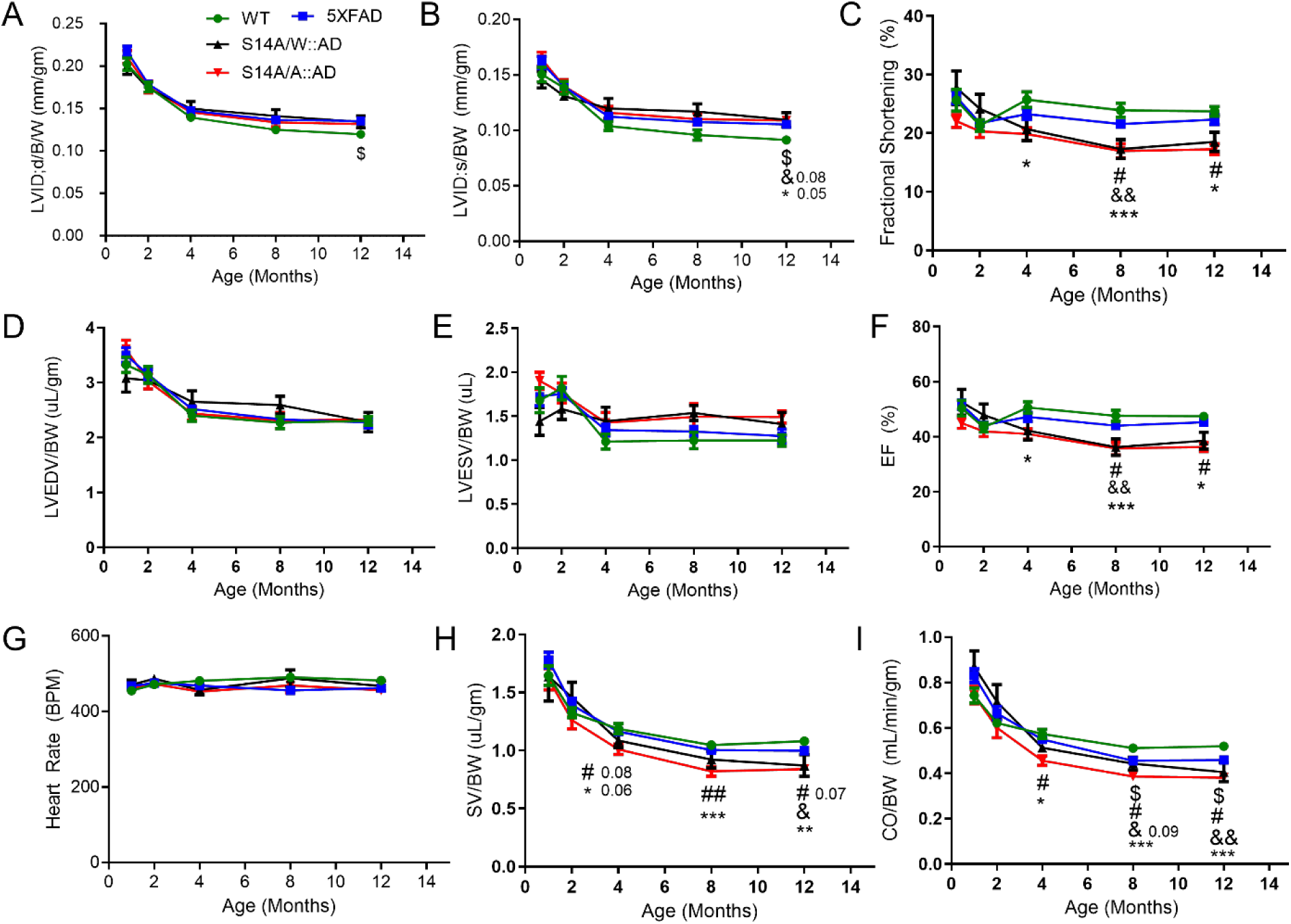
The time course of changes in the serial echocardiography-revealed cardiac functional parameters in mice with the indicated genotypes. The same cohort of mice were subjected to serial echocardiography as described in Figure 9. Mean±SEM at each time point is plotted. BW, body weight; LVID;d, left ventricular (LV) internal diameter at the end of diastole; LVID;s, LV end-systolic internal diameter; LVEDV, LV end-diastolic volume; LVSV, LV end systolic volume; EF, ejection fraction, SV, stroke volume; CO, cardiac output per minute; LVPW;d, LV posterior wall thickness at the end of diastole. 5XFAD vs. WT: $*p*<0.05; S14A/W::AD vs. WT: &p<0.05, &&p<0.01, &&&p<0.001; S14A/A::AD vs. WT: \**p*<0.05, \*\**p*<0.01, \*\*\**p*<0.001; S14A/A::AD vs. 5XFAD: #*p*<0.05, ##*p*<0.01; one-way ANOVA followed by Tukey’s tests.

**Figure 12.**
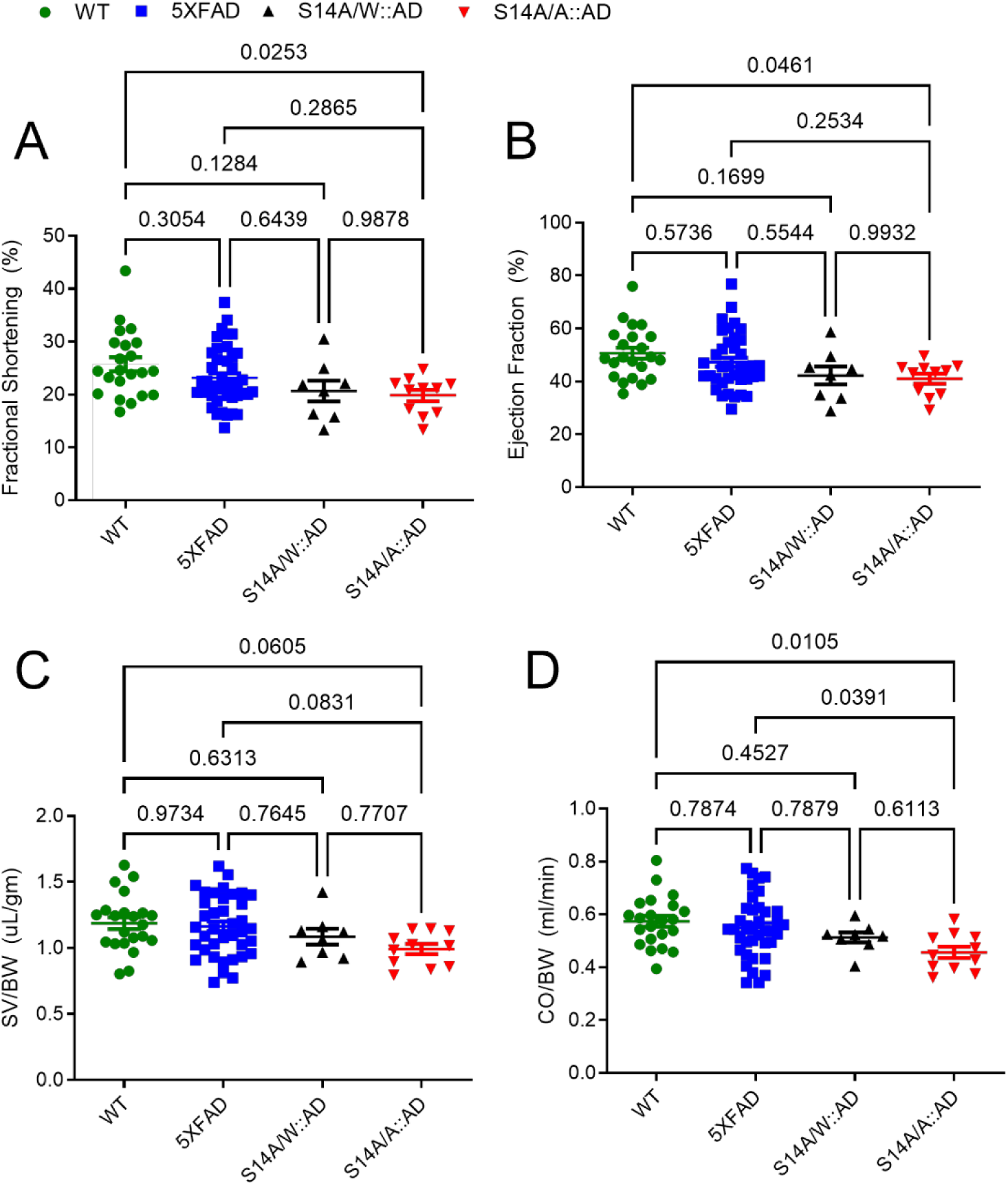
Comparison of the indicated cardiac function parameters among mice with the indicated genotypes at 4 months of age. Dot plot superimposed by mean±SEM; each dot represents an individual mouse. *P* values shown above the brackets are derived from one-way ANOVA followed by Tukey’s tests.

**Figure 13.**
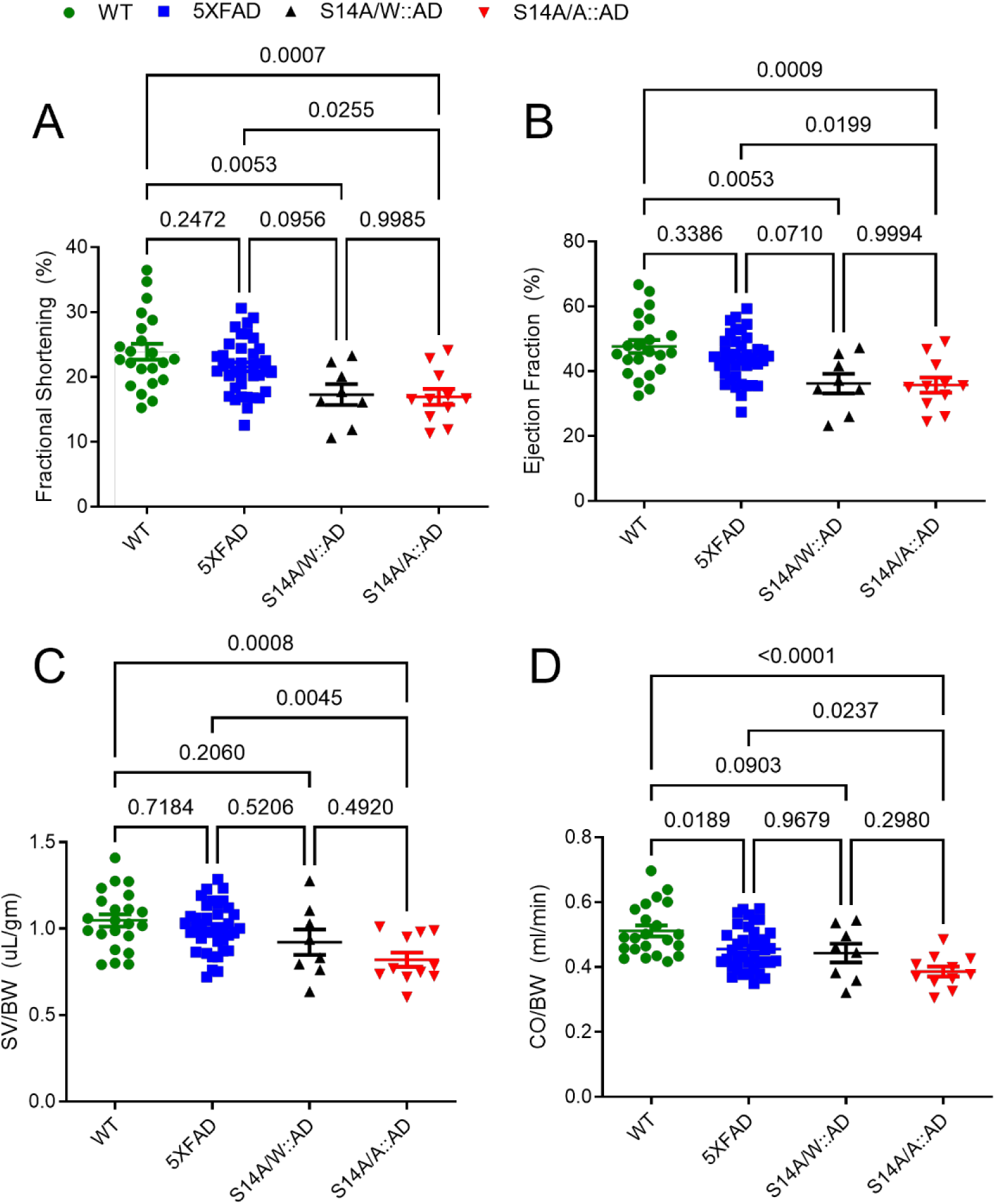
Comparison of the indicated cardiac function parameters among mice with the indicated genotypes at 8 months of age. Dot plot superimposed by mean±SEM; each dot represents an individual mouse. *P* values shown above the brackets are derived from one-way ANOVA followed by Tukey’s tests.

**Figure 14.**
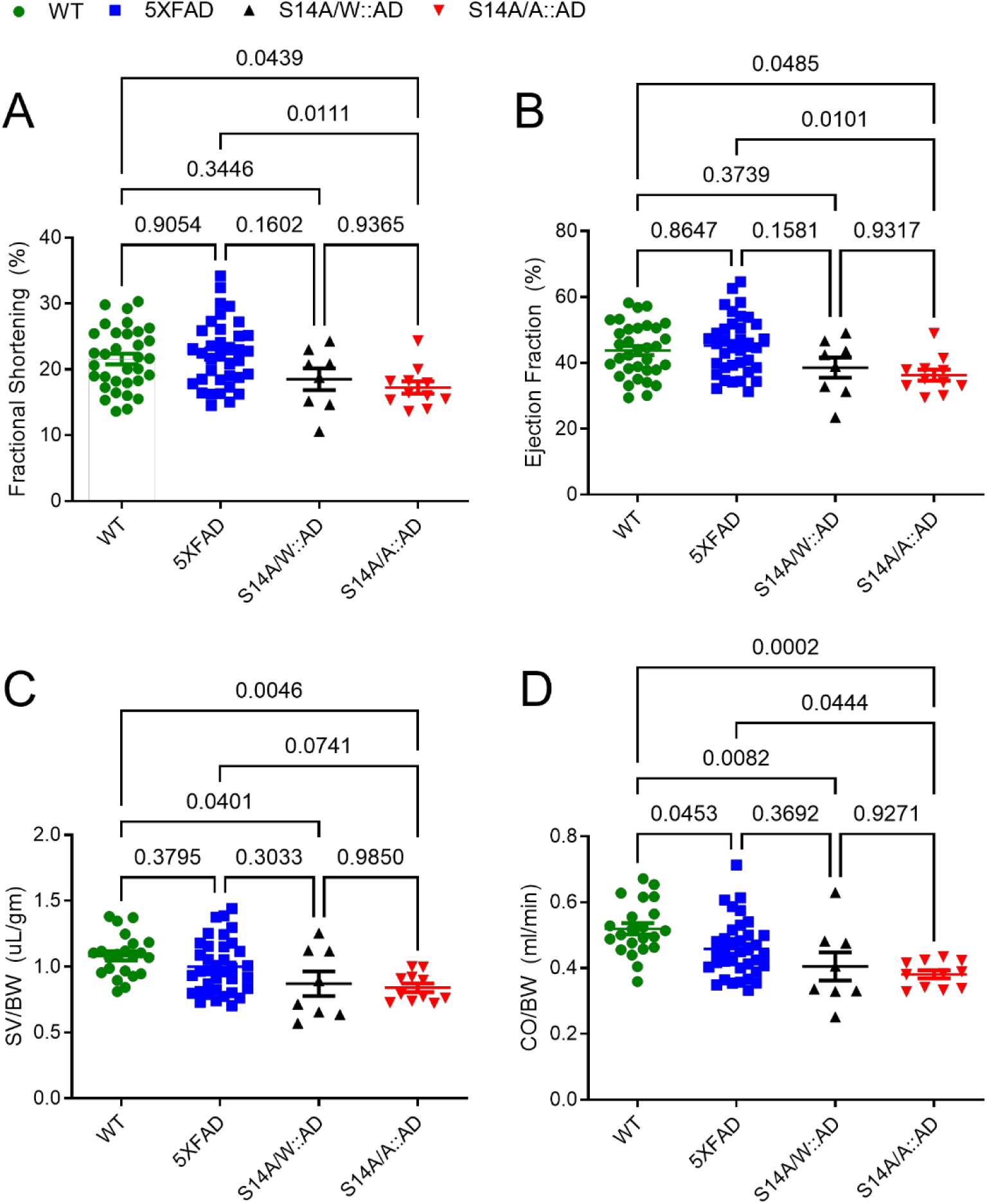
Comparison of the indicated cardiac function parameters among mice with the indicated genotypes at 12 months of age. Dot plot superimposed by mean±SEM; each dot represents an individual mouse. *P* values shown above the brackets are derived from one-way ANOVA followed by Tukey’s tests.

In conclusion, the present study establishes for the first time that homeostatic activation of 26S proteasomes by p-S14-RPN6 or by basal PKA activity protects against AD progression in both the brain and heart.

## DATA AVAILABILITY

Values for all data points in the graphs are reported in the supporting data value.xls file. Other data from this study are available upon reasonable request.

## FUNDING

This work is supported in part by NIH grants RF1AG072510, R01HL072166, and R01HL153614.

## Notes

### Competing Interest Statement

The authors have declared no competing interest.

## References

1. WHO. Available at: https://www.who.int/publications/i/item/9789240033245.

2. ADI. Available at: https://www.alzint.org/about/dementia-facts-figures/dementia-statistics/#:∼:text=There%20are%20over%2055%20million,will%20be%20in%20developing%20countries.

3. Cohen BM, Sonntag KC. Identifying the earliest-occurring clinically targetable precursors of late-onset Alzheimer’s disease. EBioMedicine 2024;106:105238.

4. Kumar A, Sidhu J, Lui F, Tsao JW. Alzheimer Disease. StatPearls. Treasure Island (FL)2024.

5. Anon. 2024 Alzheimer’s disease facts and figures. Alzheimers Dement 2024;20:3708–3821.

6. Kellogg EH, Hejab NMA, Poepsel S, Downing KH, DiMaio F, Nogales E. Near-atomic model of microtubule-tau interactions. Science 2018;360:1242–1246.

7. Glenner GG. Amyloid beta protein and the basis for Alzheimer’s disease. Prog Clin Biol Res 1989;317:857–868.

8. Glenner GG, Wong CW. Alzheimer’s disease: initial report of the purification and characterization of a novel cerebrovascular amyloid protein. Biochem Biophys Res Commun 1984;120:885–890.

9. Glenner GG, Wong CW. Alzheimer’s disease and Down’s syndrome: sharing of a unique cerebrovascular amyloid fibril protein. Biochem Biophys Res Commun 1984;122:1131–1135.

10. Baumann K, Mandelkow EM, Biernat J, Piwnica-Worms H, Mandelkow E. Abnormal Alzheimer-like phosphorylation of tau-protein by cyclin-dependent kinases cdk2 and cdk5. FEBS Lett 1993;336:417–424.

11. Alonso AC, Zaidi T, Grundke-Iqbal I, Iqbal K. Role of abnormally phosphorylated tau in the breakdown of microtubules in Alzheimer disease. Proc Natl Acad Sci U S A 1994;91:5562–5566.

12. Karran E, De Strooper B. The amyloid hypothesis in Alzheimer disease: new insights from new therapeutics. Nat Rev Drug Discov 2022;21:306–318.

13. Drummond E, Pires G, MacMurray C, et al. Phosphorylated tau interactome in the human Alzheimer’s disease brain. Brain 2020;143:2803–2817.

14. Sims JR, Zimmer JA, Evans CD, et al. Donanemab in Early Symptomatic Alzheimer Disease: The TRAILBLAZER-ALZ 2 Randomized Clinical Trial. JAMA 2023;330:512–527.

15. van Dyck CH, Swanson CJ, Aisen P, et al. Lecanemab in Early Alzheimer’s Disease. N Engl J Med 2023;388:9–21.

16. Soderberg L, Johannesson M, Nygren P, et al. Lecanemab, Aducanumab, and Gantenerumab -Binding Profiles to Different Forms of Amyloid-Beta Might Explain Efficacy and Side Effects in Clinical Trials for Alzheimer’s Disease. Neurotherapeutics 2023;20:195–206.

17. Collins GA, Goldberg AL. The Logic of the 26S Proteasome. Cell 2017;169:792–806.

18. Ciechanover A, Schwartz AL. The ubiquitin-proteasome pathway: the complexity and myriad functions of proteins death. Proc Natl Acad Sci U S A 1998;95:2727–2730.

19. Graham SH, Liu H. Life and death in the trash heap: The ubiquitin proteasome pathway and UCHL1 in brain aging, neurodegenerative disease and cerebral Ischemia. Ageing Res Rev 2017;34:30–38.

20. Haass C, Selkoe DJ. Soluble protein oligomers in neurodegeneration: lessons from the Alzheimer’s amyloid beta-peptide. Nat Rev Mol Cell Biol 2007;8:101–112.

21. Sun-Wang JL, Ivanova S, Zorzano A. The dialogue between the ubiquitin-proteasome system and autophagy: Implications in ageing. Ageing Res Rev 2020;64:101203.

22. Saez I, Vilchez D. The Mechanistic Links Between Proteasome Activity, Aging and Age-related Diseases. Curr Genomics 2014;15:38–51.

23. Schwartz AL, Ciechanover A. Targeting proteins for destruction by the ubiquitin system: implications for human pathobiology. Annu Rev Pharmacol Toxicol 2009;49:73–96.

24. Tydlacka S, Wang CE, Wang X, Li S, Li XJ. Differential activities of the ubiquitin-proteasome system in neurons versus glia may account for the preferential accumulation of misfolded proteins in neurons. J Neurosci 2008;28:13285–13295.

25. Ciechanover A, Brundin P. The ubiquitin proteasome system in neurodegenerative diseases: sometimes the chicken, sometimes the egg. Neuron 2003;40:427–446.

26. Keller JN, Hanni KB, Markesbery WR. Impaired proteasome function in Alzheimer’s disease. J Neurochem 2000;75:436–439.

27. Thibaudeau TA, Anderson RT, Smith DM. A common mechanism of proteasome impairment by neurodegenerative disease-associated oligomers. Nat Commun 2018;9:1097.

28. Russell SJ, Steger KA, Johnston SA. Subcellular localization, stoichiometry, and protein levels of 26 S proteasome subunits in yeast. J Biol Chem 1999;274:21943–21952.

29. Reits EA, Benham AM, Plougastel B, Neefjes J, Trowsdale J. Dynamics of proteasome distribution in living cells. EMBO J 1997;16:6087–6094.

30. Enenkel C, Lehmann A, Kloetzel PM. Subcellular distribution of proteasomes implicates a major location of protein degradation in the nuclear envelope-ER network in yeast. EMBO J 1998;17:6144–6154.

31. Glickman MH, Rubin DM, Coux O, et al. A subcomplex of the proteasome regulatory particle required for ubiquitin-conjugate degradation and related to the COP9-signalosome and eIF3. Cell 1998;94:615–623.

32. Wu H, Sun H, He Z, et al. The effect and mechanism of 19S proteasome PSMD11/Rpn6 subunit in D-Galactose induced mimetic aging models. Exp Cell Res 2020;394:112093.

33. Huang Q, Figueiredo-Pereira ME. Ubiquitin/proteasome pathway impairment in neurodegeneration: therapeutic implications. Apoptosis 2010;15:1292–1311.

34. Mao L, Romer I, Nebrich G, et al. Aging in mouse brain is a cell/tissue-level phenomenon exacerbated by proteasome loss. J Proteome Res 2010;9:3551–3560.

35. Gonos E. Proteasome activation as a novel anti-aging strategy. Free Radic Biol Med 2014;75 Suppl 1:S7.

36. Zhang F, Hu Y, Huang P, Toleman CA, Paterson AJ, Kudlow JE. Proteasome function is regulated by cyclic AMP-dependent protein kinase through phosphorylation of Rpt6. J Biol Chem 2007;282:22460–22471.

37. Vilchez D, Morantte I, Liu Z, et al. RPN-6 determines C. elegans longevity under proteotoxic stress conditions. Nature 2012;489:263–268.

38. Lokireddy S, Kukushkin NV, Goldberg AL. cAMP-induced phosphorylation of 26S proteasomes on Rpn6/PSMD11 enhances their activity and the degradation of misfolded proteins. Proc Natl Acad Sci U S A 2015;112:E7176–7185.

39. Lier S, Paululat A. The proteasome regulatory particle subunit Rpn6 is required for Drosophila development and interacts physically with signalosome subunit Alien/CSN2. Gene 2002;298:109–119.

40. Pathare GR, Nagy I, Bohn S, et al. The proteasomal subunit Rpn6 is a molecular clamp holding the core and regulatory subcomplexes together. Proc Natl Acad Sci U S A 2012;109:149–154.

41. Yang L, Parajuli N, Wu P, Liu J, Wang X. S14-Phosphorylated RPN6 Mediates Proteasome Activation by PKA and Alleviates Proteinopathy. Circ Res 2023;133:572–587.

42. VerPlank JJS, Lokireddy S, Zhao J, Goldberg AL. 26S Proteasomes are rapidly activated by diverse hormones and physiological states that raise cAMP and cause Rpn6 phosphorylation. Proc Natl Acad Sci U S A 2019;116:4228–4237.

43. Ahammed MS, Wang X. Promoting proteostasis by cAMP/PKA and cGMP/PKG. Trends Mol Med 2025;31:224–239.

44. Oakley H, Cole SL, Logan S, et al. Intraneuronal beta-amyloid aggregates, neurodegeneration, and neuron loss in transgenic mice with five familial Alzheimer’s disease mutations: potential factors in amyloid plaque formation. J Neurosci 2006;26:10129–10140.

45. Yang L, Ahammed MS, Wu P, Sternburg JO, Liu J, Wang X. Genetic blockade of the activation of 26S proteasomes by PKA is well tolerated by mice at baseline. Am J Cardiovasc Dis 2024;14:90–105.

46. Leger M, Quiedeville A, Bouet V, et al. Object recognition test in mice. Nat Protoc 2013;8:2531–2537.

47. Lueptow LM. Novel Object Recognition Test for the Investigation of Learning and Memory in Mice. J Vis Exp 2017.

48. Park CR, Zoladz PR, Conrad CD, Fleshner M, Diamond DM. Acute predator stress impairs the consolidation and retrieval of hippocampus-dependent memory in male and female rats. Learn Mem 2008;15:271–280.

49. Alamed J, Wilcock DM, Diamond DM, Gordon MN, Morgan D. Two-day radial-arm water maze learning and memory task; robust resolution of amyloid-related memory deficits in transgenic mice. Nat Protoc 2006;1:1671–1679.

50. Adegoke OO, Qiao F, Liu Y, Longley K, Feng S, Wang H. Overexpression of Ubiquilin-1 Alleviates Alzheimer’s Disease-Caused Cognitive and Motor Deficits and Reduces Amyloid-beta Accumulation in Mice. J Alzheimers Dis 2017;59:575–590.

51. Zhang H, Pan B, Wu P, et al. PDE1 inhibition facilitates proteasomal degradation of misfolded proteins and protects against cardiac proteinopathy. Sci Adv 2019;5:eaaw5870.

52. Reyes-Marin KE, Nunez A. Seizure susceptibility in the APP/PS1 mouse model of Alzheimer’s disease and relationship with amyloid beta plaques. Brain Res 2017;1677:93–100.

53. Barbour AJ, Gourmaud S, Lancaster E, et al. Seizures exacerbate excitatory: inhibitory imbalance in Alzheimer’s disease and 5XFAD mice. Brain 2024;147:2169–2184.

54. Troncone L, Luciani M, Coggins M, et al. Abeta Amyloid Pathology Affects the Hearts of Patients With Alzheimer’s Disease: Mind the Heart. J Am Coll Cardiol 2016;68:2395–2407.

55. Murphy J, Le TNV, Fedorova J, et al. The Cardiac Dysfunction Caused by Metabolic Alterations in Alzheimer’s Disease. Front Cardiovasc Med 2022;9:850538.

56. Tublin JM, Adelstein JM, Del Monte F, Combs CK, Wold LE. Getting to the Heart of Alzheimer Disease. Circ Res 2019;124:142–149.

57. Cermakova P, Eriksdotter M, Lund LH, Winblad B, Religa P, Religa D. Heart failure and Alzheimer’s disease. J Intern Med 2015;277:406–425.

58. Jagust W. Untangling vascular dementia. Lancet 2001;358:2097–2098.

59. Alosco ML, Hayes SM. Structural brain alterations in heart failure: a review of the literature and implications for risk of Alzheimer’s disease. Heart Fail Rev 2015;20:561–571.

60. Li D, Parks SB, Kushner JD, et al. Mutations of presenilin genes in dilated cardiomyopathy and heart failure. Am J Hum Genet 2006;79:1030–1039.

